# *Plasmodium falciparum* hemozoin-associated biomolecules induce brain endothelial cell barrier disruption in an in vitro model of cerebral malaria

**DOI:** 10.64898/2026.03.12.711413

**Authors:** Kelly A. Crotty, Ioana Clotea, Beatrix Ueberheide, Michael Cammer, Joseph Sall, Alice Liang, Ana Rodriguez

## Abstract

Cerebral malaria is a major complication of *Plasmodium falciparum* infection that occurs upon the sequestration of infected red blood cells (iRBCs) in brain capillaries, resulting in the loss of endothelial barrier integrity, brain swelling, and frequently long-term sequelae or death. *P. falciparum*-iRBCs cause the disruption of human brain microvascular endothelial cell barrier integrity in vitro, mimicking the microenvironment of cerebral malaria, yet the specific mechanisms mediating this process remain unknown. Upon infection of the host RBCs, *P. falciparum* produces hemozoin, a crystal form of heme generated following the degradation of hemoglobin by the parasite. Here we show that the endothelial barrier-disrupting activity is found entirely in the hemozoin fraction of *P. falciparum*-iRBCs. This activity is not caused by the hemozoin crystal itself, which is not able to induce barrier disruption, but by the biomolecules that are associated with it. Treatment of purified *P. falciparum* hemozoin with proteases inhibits the disruption of endothelial barrier integrity caused by the hemozoin, indicating an important role for proteins in the disruption of the barrier. Conversely, treatment with nucleases did not affect hemozoin barrier disrupting activity. These results identify a key molecular mechanism in the *P. falciparum*-mediated brain endothelial barrier disruption during cerebral malaria and may open new avenues for the treatment of this complication.

**IMPORTANCE:** While several specific biomolecules have been proposed to contribute to the disruption of endothelial barrier integrity in cerebral malaria, no single *P. falciparum-* or host-derived factor has been definitively identified as the primary driver of this disruption. Here, we identify the brain endothelial barrier-disruptive *P. falciparum-*iRBC-derived activity to be caused by biomolecules bound to hemozoin, identifying a key, novel mechanism in the pathogenesis of cerebral malaria. The finding that *P. falciparum* hemozoin also disrupts a pulmonary endothelial cell barrier opens the possibility that this mechanism underlies other severe malaria complications. The implication of *P. falciparum-*iRBC-derived proteins in this mechanism is in line with previous reports, providing a novel interpretation of these findings in the context of hemozoin-binding. This knowledge provides a new perspective in the search for specific molecules and mechanisms involved in barrier disruption, which may lead to the development of much-needed specific treatments for cerebral malaria.

## INTRODUCTION

Malaria remains one of the top causes of death in low-income countries, with 263 million cases and 597,000 deaths in 2023 alone.^1–4^ A major contributor to this high number of deaths is cerebral malaria (CM), a life-threatening complication of *Plasmodium falciparum* infection that often presents with seizures and delirium, leading to coma.^5^ CM is frequently fatal with 20% of CM patients treated with antimalarials still dying from the disease.^6^ Of those who are treated and survive, up to 25% will have long-lasting neurological or cognitive sequelae.^5^ Despite the critical nature of this complication, there is currently no drug to specifically treat CM.

The pathogenesis of CM occurs in the microvasculature of the brain, at the forefront of the blood-brain barrier (BBB), where *P. falciparum*-infected red blood cells (iRBCs) bind to the surface of the endothelial cells lining the BBB.^7–10^ This binding leads to the blockage of the vessel following the clumping or agglutination of iRBCs with other iRBCs, RBCs, or platelets.^11–13^ As the iRBCs complete the erythrocytic cycle in these sequestered areas, they rupture and release their contents, creating a microenvironment with a high concentration of iRBC molecules in close proximity to the brain endothelial cells. Previous reports have shown the importance of this rupture step in in vitro models of CM as intracellular components of the iRBCs disrupt the inter-endothelial junctions, resulting in a decrease in barrier integrity.^14–22^ Once the endothelial junctions are compromised, fluid leaks through the BBB, causing brain swelling by way of vasogenic edema, which is highly correlated with death in CM.^23,24^ While several specific *P. falciparum*-derived molecules have been proposed to contribute to the decrease in barrier integrity, such as histidine rich protein II (HRPII),^16–18^ histones,^19^ and glycophosphatidylinositols,^20,21^ no single iRBC-derived factor has been definitively identified as the primary driver of BBB breakdown.

Hemozoin (Hz) is an insoluble crystal form of heme that is formed as a byproduct of heme detoxification as the *Plasmodium* parasite develops within the RBC.^25–27^ Hz is amphiphilic, meaning that one face is polar and hydrophilic, and the other is hydrophobic and lipophilic, resulting in a wide biomolecule binding capacity that includes nucleic acids, proteins, and lipids.^25,28^ As the *P. falciparum-*iRBCs complete their life cycle, they rupture and release Hz,^29^ and therefore the bound biomolecules, which can be found in close proximity to the endothelial cells lining the BBB.^30^ It has previously been demonstrated that lowering the levels of Hz through the inhibition of the heme pathway, either by drugs or genetic knockout, results in less aggressive CM symptoms in an experimental CM mouse model.^31,32^ This mouse study points to a role for Hz in the pathogenesis of CM,^31^ however the molecular mechanism underlying this effect and whether this is applicable to human CM is still unclear.

In this study, we identified that the endothelial-barrier disruptive activity of *P. falciparum*-iRBCs is entirely caused by biomolecules bound to *P. falciparum* Hz, but not the Hz crystal itself. Additionally, treatment of *P. falciparum* Hz with proteases inhibits the barrier disruption caused by the hemozoin, which is in agreement with previous reports identifying specific proteins as mediators of barrier disruption.^16–19^ Further elucidating the mechanism behind brain endothelial barrier disruption in CM may lead to the development of targeted therapeutics which could drive down the high morbidity and mortality rates of this complication.

## RESULTS

### Hemozoin purified from *P. falciparum* schizont-stage iRBCs disrupts the barrier integrity of immortalized human brain microvascular endothelial cells

Previous reports have demonstrated that *P. falciparum*-iRBCs disrupt the barrier integrity of primary and immortalized human brain microvascular endothelial cells (HBMECs) upon their rupture and release of intracellular contents,^14,33,34^ and that a lysate of schizont-stage iRBCs has similar activity.^33^ Here we observed that when *P. falciparum* (3D7 strain) iRBCs at the end of the erythrocytic cycle are either lysed by freeze-thawing (iRBC lysate) or allowed to naturally rupture (naturally ruptured iRBCs) and the collected materials are then added to HBMECs, they similarly induce barrier disruption (**Figure 1A**). This disruption of HBMEC barrier integrity was also induced by iRBC lysate and naturally ruptured iRBCs from the Dd2, NF54, and W2 strains of *P. falciparum* (**Figures S1-2**).

**Figure 1:**
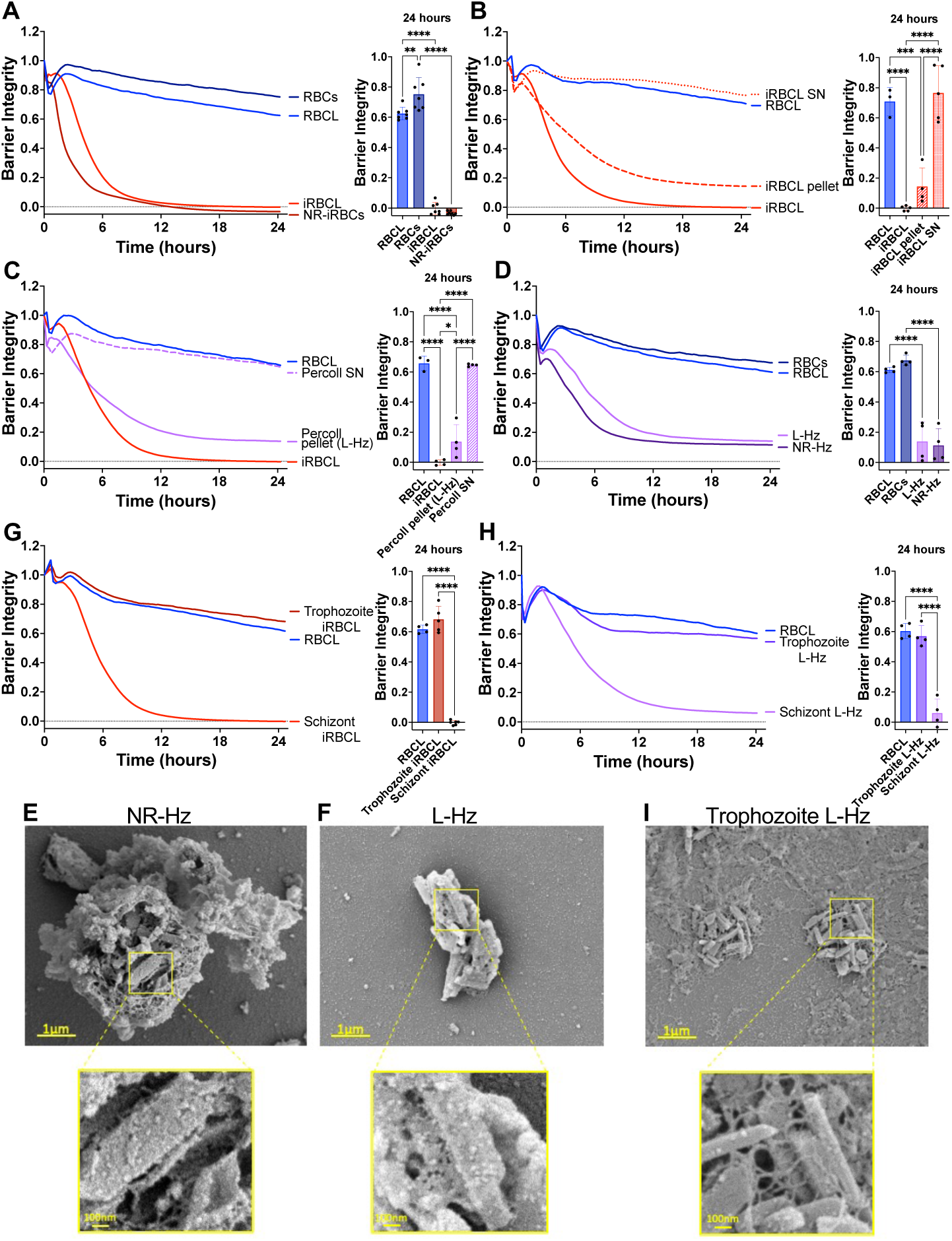
Hz purified from *P. falciparum* schizont-stage iRBCs disrupts the HBMEC barrier. RBCs (blue lines) or iRBCs (red lines) were incubated at 37°C overnight to allow iRBC to rupture naturally (RBCs, NR-iRBCs) or were lysed by freeze-thaw (RBCL, iRBCL) before adding to a monolayer of HBMECs. Barrier integrity of HBMEC monolayers was measured by xCELLigence as impedance every 15min over 24h. Bars represent barrier integrity at 24h (**A-D, G-H**). (**B**) iRBCL was spun down and separated into the pellet and supernatant (SN) fractions before adding to HBMECs. (**C-D, G-H**) Schizont iRBCL (iRBCL), NR-iRBCs, or trophozoite iRBCL were spun over a 10% Percoll solution to obtain pellet (L-Hz from iRBCL or NR-Hz from NR-iRBCs) and/or SN. (**C-D**) Purified Hz concentration matched to measured Hz in iRBCs. (**H**) Purified Hz concentration is 18.88µg/cm^2^. Results represent the average from 2+ independent experiments performed in duplicate with standard deviations. Statistical significance was determined by one-way ANOVA with Tukey’s multiple comparisons test where * = *P* < 0.05, ** = *P* < 0.01, *** = *P* < 0.001, and **** = *P* < 0.0001. Representative images of (**E**) NR-Hz, (**F**) L-Hz, or (**I**) trophozoite L-Hz fixed in the presence of malachite green and imaged under SEM.

To identify the active component within the iRBC lysate, we centrifuged the iRBC lysate to generate pellet and supernatant fractions, observing that the HBMEC barrier-disruptive activity was restricted to the pellet fraction (**Figure 1B**). We further separated the iRBC lysate using density centrifugation (10% Percoll) to separate Hz from the rest of the iRBC lysate components.^35,36^ The Percoll centrifugation pellet, composed almost exclusively of *P. falciparum* Hz, retained the barrier-disruptive activity, while the supernatant fraction did not (**Figure 1C**). The disruption caused by the purified Hz, but not the supernatant fraction, was confirmed in a parallel filter permeability assay (**Figure S3A**).

We then verified whether the disruptive activity of Hz purified from iRBC lysate was also found in Hz purified from naturally ruptured iRBCs, finding that they both exhibited similar endothelial barrier disruptive activity (**Figures 1D, S4-5, and S6A**). Despite similar activity, Hz obtained from these two different preparations of *P. falciparum*-iRBCs showed different associated components, as the Hz from the naturally ruptured iRBCs remains enclosed within the food vacuole (**Figure 1E**),^37^ while the freeze-thawing process used to generate the iRBC lysate releases free Hz crystals (**Figure 1F**). The observed disruption of the endothelial barrier by Hz from naturally ruptured iRBCs is consistent with previous reports of purified food vacuoles inducing endothelial disruption.^20^

We also observed that only the schizont-stage iRBC lysate and Hz purified from it exhibit barrier-disrupting activity, while the trophozoite-stage counterparts do not (**Figures 1G-I and S6B**), indicating that endothelial barrier disruption can be attributed to Hz derived specifically from schizont-stage iRBCs.

### *P. falciparum* hemozoin disrupts the barrier of other endothelial cell types

We also observed that Percoll-purified, schizont-stage Hz from iRBC lysate disrupted the barrier integrity of primary HBMECs, primary human cardiac microvascular endothelial cells, and immortalized human pulmonary microvascular endothelial cells (**Figure 2**). These results indicate that the barrier disrupting activity of Hz is not restricted to the brain endothelium and may have implications for pathogenesis in different organs where iRBCs sequester during *P. falciparum* infection.^38^

**Figure 2:**
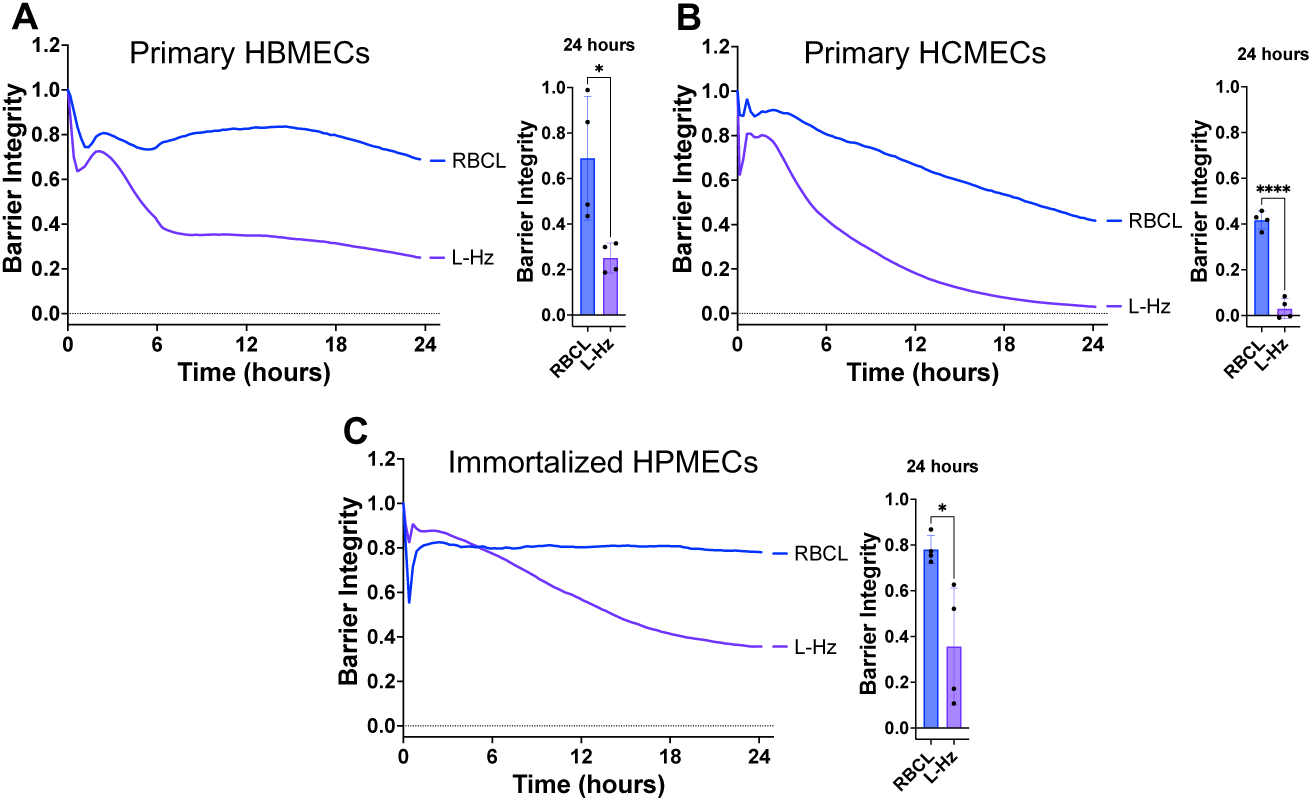
*P. falciparum* Hz disrupts the barrier of other endothelial cells. RBC lysate (RBCL) or Hz from iRBC lysate (L-Hz; 18.88µg/cm^2^) were added to a monolayer of primary human brain microvascular endothelial cells (HBMECs, **A**), primary human cardiac microvascular endothelial cells (HCMECs, **B**), or immortalized human pulmonary microvascular endothelial cells (HPMECs, **C**). Barrier integrity of HBMEC monolayers was measured by xCELLigence as impedance every 15min over 24h. Bars represent barrier integrity at 24h. Results represent the average from 2 independent experiments performed in duplicate with standard deviations. Statistical significance was determined by unpaired t test where * = *P* < 0.05 and **** = *P* < 0.0001.

### Hemozoin purified from *P. falciparum*-iRBC lysate, but not synthetic hemozoin, disrupts the brain endothelial barrier in a dose-dependent manner

As an orthogonal approach, we purified the Hz from iRBC lysate using a strong magnet, as Hz is paramagnetic.^39^ We observed that Hz purified using either Percoll centrifugation (**Figure 3A**) or magnetic separation (**Figure 3B**) disrupts the HBMEC barrier integrity in a similar dose-dependent manner, which can also be visualized by fluorescence microscopy (**Figure 3C**). However, when commercially available synthetic Hz was added to the HBMECs at the same concentrations as the Hz purified from iRBC lysate in Figures 3A-B, there was minimal disruption of the endothelial barrier (**Figure 3D**), as previously reported.^20^ This lack of disruption was also observed in a parallel filter permeability assay where the highest dose of synthetic Hz from Figure 3D did not induce significant permeability in the HBMEC monolayer after 24h of incubation (**Figure S3B**).

**Figure 3:**
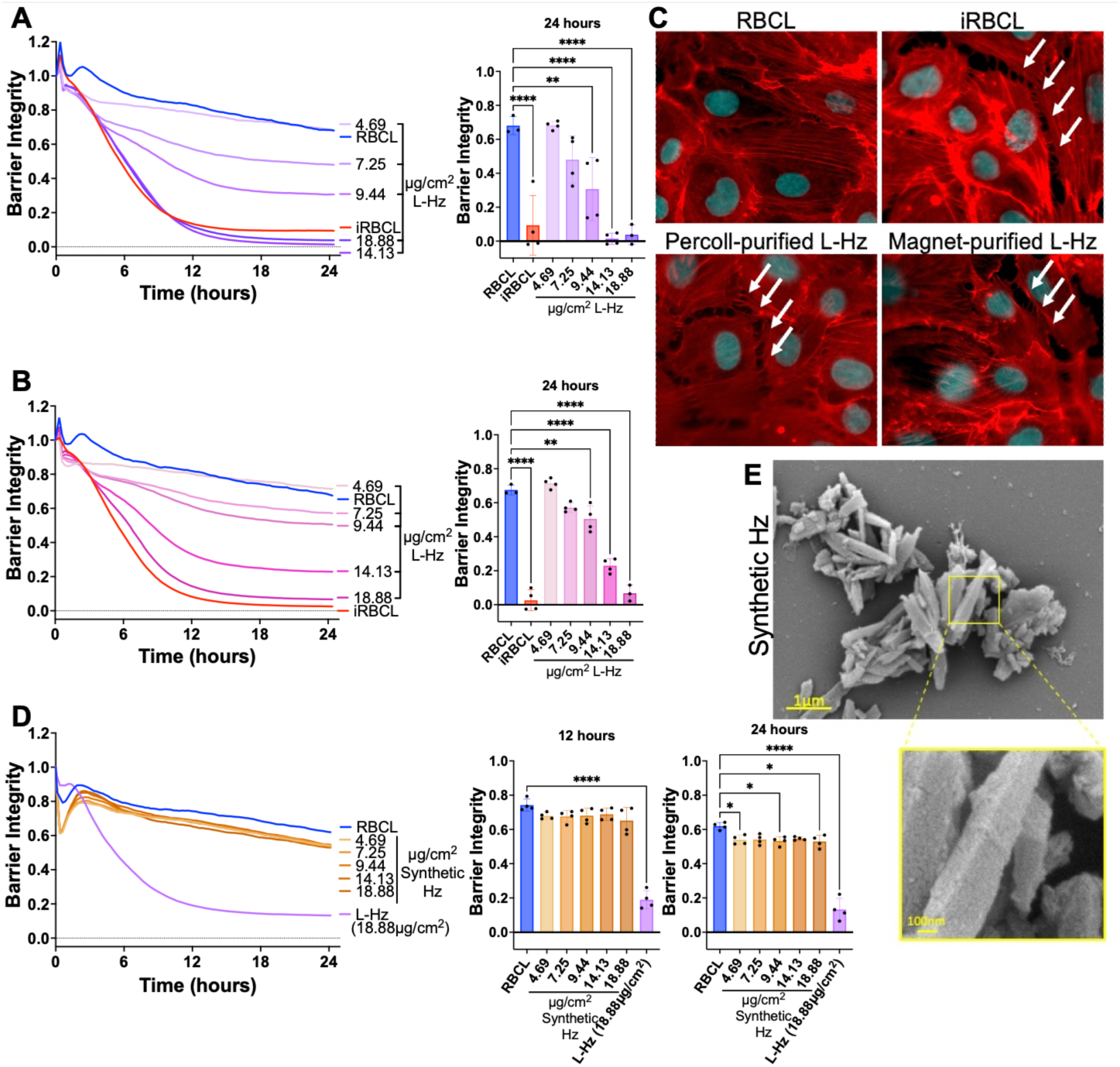
*P. falciparum* Hz, but not synthetic Hz, disrupts the HBMEC barrier in a dose-dependent manner. Different concentrations of Percoll-purified (**A**) or magnet-purified (**B**) Hz from iRBC lysate (L-Hz) were added to a monolayer of HBMECs along with RBC lysate (RBCL) and iRBC lysate (iRBCL). The measured Hz concentration of the iRBCL ranged from 7.19 –9.25µg/cm^2^ in the independent experiments. (**C**) Representative images of monolayers of HBMECs that were incubated for 4h with the indicated stimuli (L-Hz at 18.88µg/cm^2^) before fixation and actin (Phalloidin; red) and nuclei (Hoechst; cyan) staining. (**D**) Synthetic Hz at the same concentrations as the L-Hz in A-B, RBCL, and L-Hz were added to a monolayer of HBMECs. (**A-B, D**) Barrier integrity of HBMEC monolayers was measured by xCELLigence as impedance every 15min over 24h. Bars represent barrier integrity at 24h. Results represent the average from 2 independent experiments performed in duplicate with standard deviations. Statistical significance was determined by one-way ANOVA with Tukey’s multiple comparisons test where * = *P* < 0.05, ** = *P* < 0.01, and **** = *P* < 0.0001. (**E**) Representative images of synthetic Hz fixed in the presence of malachite green and imaged under SEM.

A major difference between *P. falciparum*-derived and synthetic Hz is that the parasite Hz has a diverse range of iRBC biomolecules bound to it, such as nucleic acids, proteins, and lipids.^25,28^ This is also apparent in scanning electron microscopy (SEM) images where Hz from *P. falciparum-*iRBCs presents an uneven surface, likely reflecting the large quantities of biomolecules bound to it (**Figure 1E-F**), whereas synthetic Hz shows the even, angular appearance of a bare crystal (**Figure 3E**).

We also observed that the barrier-disrupting activity of the iRBC lysate is sensitive to physical and chemical changes, such as heating at 60°C or 95°C, sonication, and high or low pH (**Figure S7**). Taken together, these results indicate that the biomolecules bound to *P. falciparum* Hz are responsible for the HBMEC barrier disruption activity, rather than the Hz crystal itself.

### Hemozoin internalization by brain endothelial cells is not required for barrier disruption

To investigate mechanisms for *P. falciparum*-derived Hz disruption of the HBMEC barrier integrity, we first determined whether Hz internalization by the HBMECs is required for barrier disruption. We observed that the Hz present in the iRBC lysate and Hz purified from it are not internalized by the HBMECs, while synthetic Hz is internalized, colocalizing with a lysosomal marker (LAMP1) (**Figure 4**). These results show that while HBMECs can internalize the bare synthetic Hz crystal, they do not internalize *P. falciparum*-derived Hz, indicating that Hz internalization is not required for the disruption of HBMEC barrier integrity.

**Figure 4:**
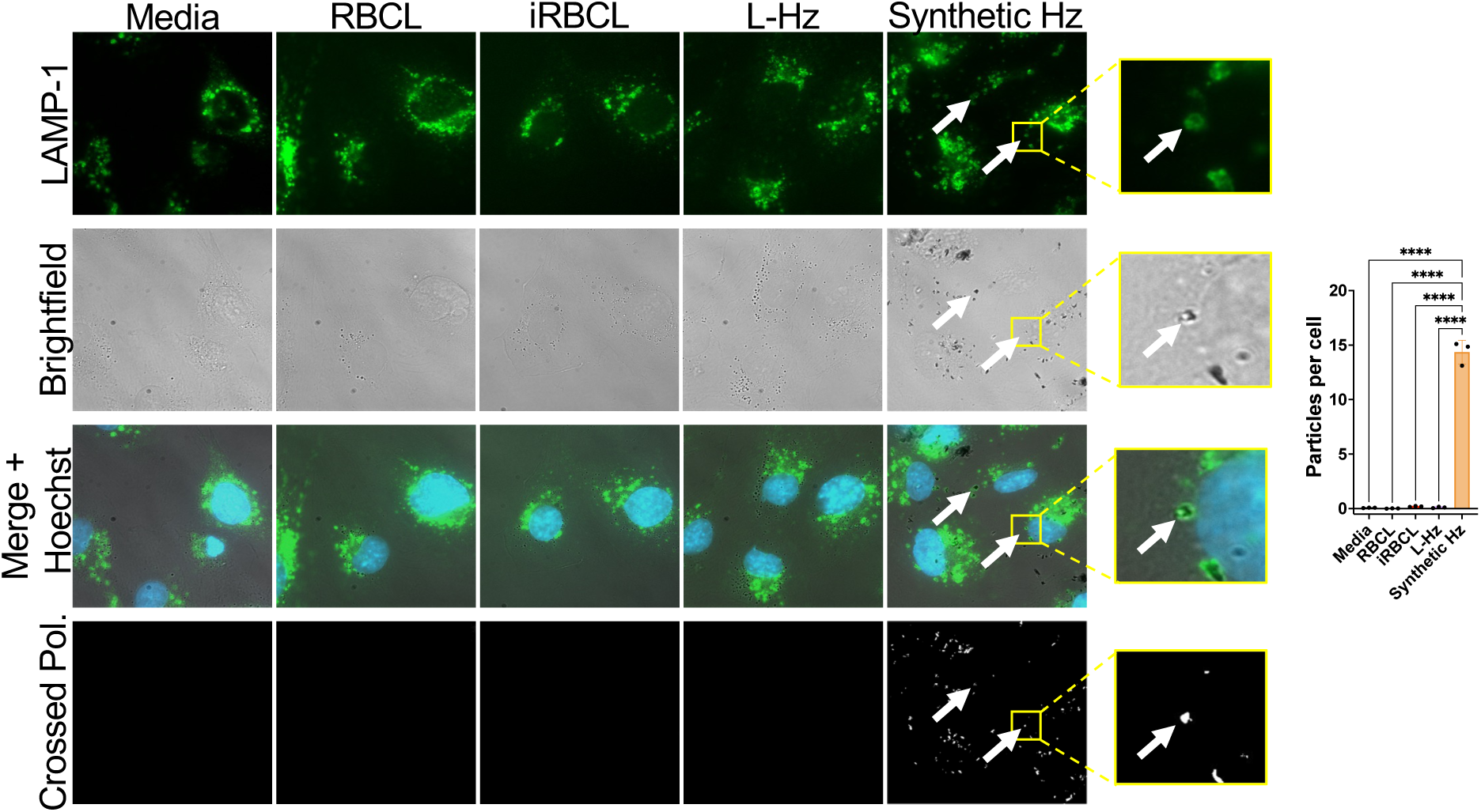
*P. falciparum* Hz is not internalized by HBMECs. Representative images of monolayers of HBMECs that were incubated for 2h with RBC lysate (RBCL), iRBC lysate (iRBCL), Hz from iRBCL (L-Hz) or synthetic Hz (concentrations matched to Hz in iRBCL) before being washed, fixed, and stained for lysosomes (LAMP-1; green) and nuclei (Hoechst; blue). Crossed polarization (Crossed Pol.) was used to illuminate crystal-like structures. White arrows show examples of lysosomal LAMP1 staining surrounding a hemozoin crystal. For quantification, 100 cells in 3 different wells per condition were counted along with the number of crystal-like particles illuminated by crossed polarization. Statistical significance was determined by one-way ANOVA with Tukey’s multiple comparisons test where **** = *P* < 0.0001.

### Nucleic acids do not contribute to the disruption of HBMEC barrier by *P. falciparum* iRBCs

We explored the contribution of nucleic acids to the disruption of endothelial barrier integrity by treating the iRBC lysate with DNase (**Figure 5A-B**) or RNase (**Figure 5C-D**). We did not observe any attenuation of the iRBC lysate activity following a strong reduction in DNA or RNA, indicating that nucleic acids do not contribute to the HBMEC barrier disruption.

**Figure 5:**
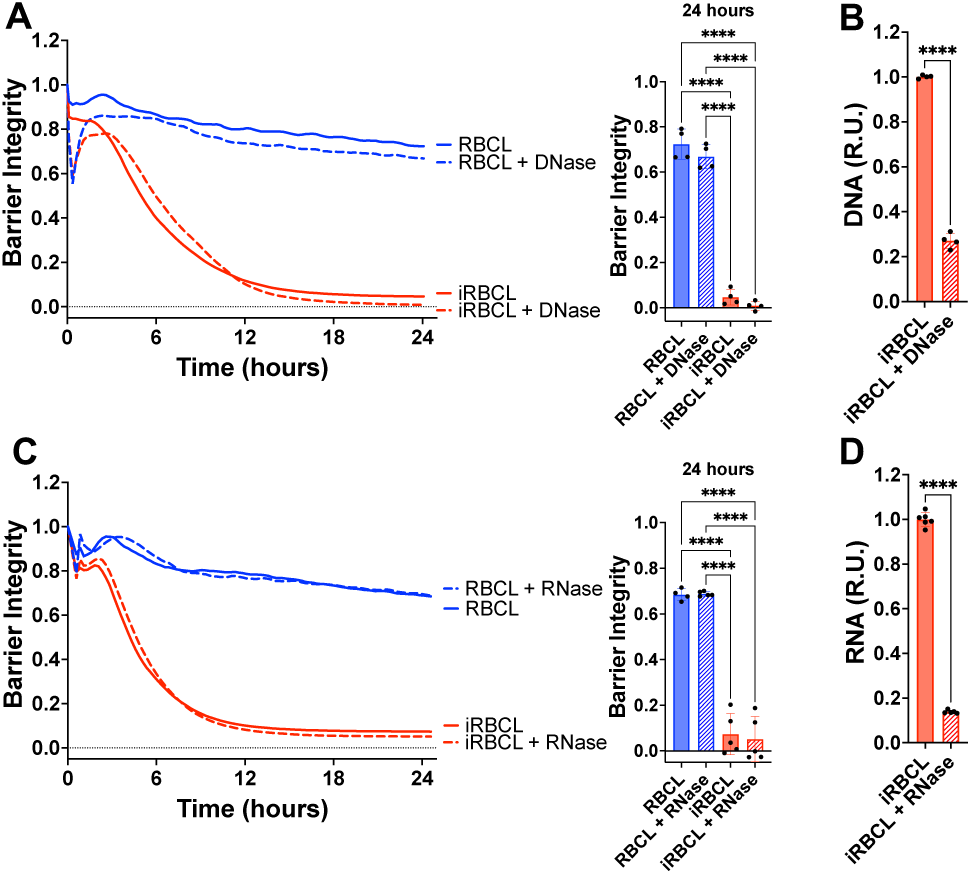
Nucleic acids do not contribute to the disruption of HBMEC barrier by *P. falciparum* iRBCs. RBC lysate (RBCL) or iRBC lysate (iRBCL) were treated with either DNase (**A**) or RNase (**C**) for 30min prior to adding to a monolayer of HBMECs. Barrier integrity of HBMEC monolayers was measured by xCELLigence as impedance every 15min over 24h. Bars represent barrier integrity at 24h. (**B**) DNA and (**D**) RNA concentrations were determined by SYBR Green and SYTO RNASelect, respectively. The data was normalized to the iRBCL and is represented in relative units (R.U.). Results represent the average from 2+ independent experiments performed in duplicate with standard deviations. Statistical significance was determined by one-way ANOVA with Tukey’s multiple comparisons test (**A, C**) or unpaired t test (**B, D**) where *** = *P* < 0.001 and **** = *P* < 0.0001.

### Proteases inhibit the brain endothelial barrier-disrupting activity of *P. falciparum*-iRBC lysate

We next investigated the role of Hz-associated proteins by treating the iRBC lysate and Hz purified from it with trypsin or Proteinase K prior to incubation with the HBMECs, finding a strong decrease in barrier disrupting activity (**Figures 6A-D**), which indicates that Hz-bound proteins play an important role in HBMEC barrier disruption.

**Figure 6:**
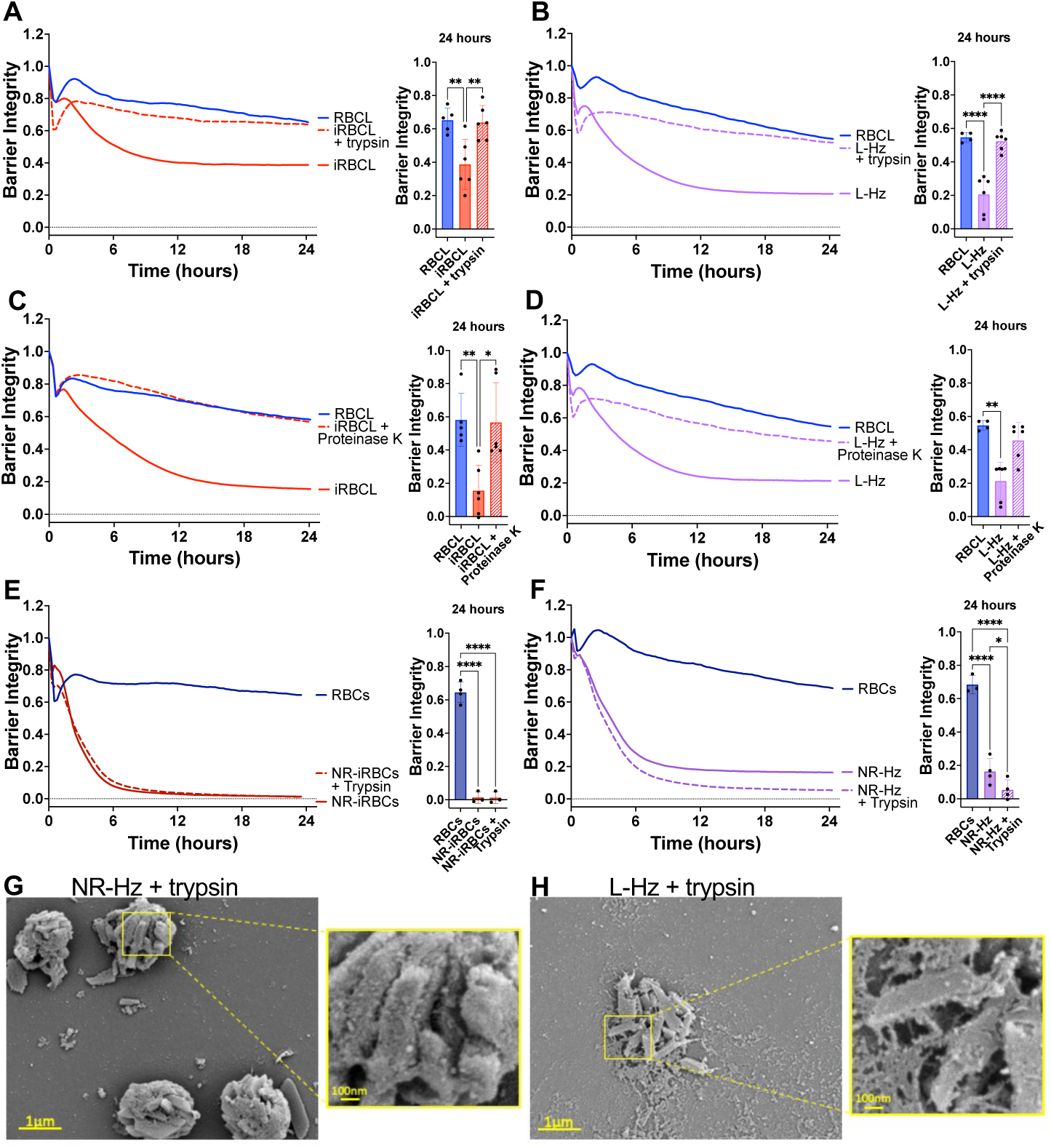
Proteases inhibit endothelial barrier disruption caused by iRBC lysate and its purified Hz. Lysed (iRBCL, **A**) or naturally ruptured (NR-iRBCs, **E**) iRBCs or Hz from iRBCL (L-Hz, **B**) or NR-iRBCs (NR-Hz, **F**) (concentration matched to Hz in iRBCL) were treated with trypsin for 30min prior to adding to a monolayer of HBMECs along with RBC lysate (RBCL) or RBCs. iRBCL (**C**) or L-Hz (**D**; concentration matched to Hz in iRBCL) were treated with Proteinase K for 30min prior to washing out the enzyme and adding to a monolayer of HBMECs along with RBCL. Barrier integrity of HBMEC monolayers was measured by xCELLigence as impedance every 15min over 24h. Bars represent barrier integrity at 24h. Results represent the average from 2+ independent experiments performed in duplicate with standard deviations. Statistical significance was determined by one-way ANOVA with Tukey’s multiple comparisons test (**A-B, E-F**) or Kruskal-Wallis test with Dunn’s multiple comparisons test (**C-D**) where * = *P* < 0.05, ** = *P* < 0.01, and **** = *P* < 0.0001. Representative images of trypsin-treated (**G**) NR-Hz or (**H**) L-Hz fixed in the presence of malachite green and imaged under SEM.

However, we also observed that the barrier-disrupting activity of naturally ruptured iRBCs or Hz purified from them was not inhibited by trypsin treatment (**Figures 6E-F**). Since Hz is contained within the food vacuole of the iRBCs,^40^ it is likely that the protease did not have access to Hz-bound proteins in the naturally ruptured iRBCs and the Hz purified from them. This is also suggested by the SEM images of Hz from naturally ruptured iRBCs, where the surrounding membrane of the food vacuole is visible in both the untreated (**Figure 1E**) and trypsin-treated preparations (**Figure 6G**). Conversely, trypsin-treated Hz purified from the iRBC lysate appears to be more ‘naked’ (**Figure 6H**) than its untreated counterpart (**Figure 1F**) suggesting that trypsin has access to the proteins bound to the free Hz crystals.

### Identification of *P. falciparum-*iRBC proteins bound to hemozoin

Electrophoresis separation revealed a considerable number of proteins bound to Hz purified from the iRBC lysate, though fewer than observed in the whole iRBC lysate (**Figure S8**). Mass spectrometric proteomic analysis of Hz-associated proteins identified 1,890 *P. falciparum* and 196 human proteins (**Table S1**). Comparison of this dataset to a comprehensive *P. falciparum*-iRBC proteomic dataset identified that approximately 72% of all *P. falciparum* proteins expressed at the schizont stage are able to bind to Hz.^41^ We observed that the top 10 most abundant *P. falciparum* Hz-bound proteins (**Table 1**), are found within the top 6% most abundant *P. falciparum* proteins in the whole iRBC,^41^ suggesting a lack of specificity in the protein binding to Hz. However, it is important to note that several *P. falciparum* histones, which have been implicated in the endothelial barrier disruption in a model of CM,^42^ are found among the most abundant *P. falciparum* proteins bound to Hz (**Table 1**).

**Table 1:** Top 10 most abundant *P. falciparum* proteins associated with Hz purified from iRBC lysate (L-Hz) ranked by relative abundance. Columns show protein codes, gene IDs, and rank by intensity in schizont-stage *P. falciparum* proteins in DDA dataset from Siddiqui et al. (2020).^41^.

| UniProt Code | PlasmoDB ID | Rank in whole iRBC<sup>41</sup> | Protein Name |
| --- | --- | --- | --- |
| Q8I0U8 | PF3D7_0930300 | 1 | Merozoite surface protein 1 |
| Q8IIV2 | PF3D7_1105000 | 14 | Histone H4 |
| C0H571 | PF3D7_0929400 | 24 | High molecular weight rhoptry protein 2 |
| Q8ILZ1 | PF3D7_1410400 | 22 | Rhoptry-associated protein 1 |
| Q8I492 | PF3D7_0500800 | 10 | Mature parasite-infected erythrocyte surface antigen |
| Q8I4R5 | PF3D7_1252100 | 144 | Rhoptry neck protein 3 |
| Q8IIV1 | PF3D7_1105100 | 29 | Histone H2B |
| O77310 | PF3D7_0302500 | 79 | Cytoadherence linked asexual protein 3.1 |
| O97320 | PF3D7_0320900 | 31 | Histone H2A |
| C6KT18 | PF3D7_0617800 | 141 | Histone H2A |

## DISCUSSION

Disruption of the BBB integrity is a fundamental process in the pathogenesis of CM, leading to brain swelling and frequently death,^23^ yet the mechanism underlying this process remains poorly understood.^43^ Disruption of the BBB in CM occurs following the binding of *P. falciparum*-iRBCs to brain endothelial cells, causing sequestration and impeded blood flow in the area,^44^ resulting in both vasogenic and hypoxia-associated cytotoxic edema in distinct parts of the brain.^24^ Previous studies using 2D and 3D models have established that *P. falciparum*-iRBCs disrupt brain endothelial cell barrier integrity in vitro,^14,22,33,34,45^ providing useful tools to identify the molecular components mediating this disruption.^16–21^ It has been observed that several parasite-derived biomolecules can mediate endothelial barrier disruption.^16–21^ Despite these findings, no definitive iRBC factor has been characterized as the driving force behind CM, which is likely a multifactorial phenomenon.^46^

This study has identified that the iRBC molecules responsible for the endothelial barrier disrupting activity are bound to the Hz of mature schizont iRBCs, while no activity was observed in other fractions. In the context of CM, where iRBCs are sequestered in brain capillaries, the release of Hz by naturally rupturing iRBCs at the end of the erythrocytic cycle would deliver Hz directly over the endothelial cells in a microenvironment of impeded blood flow.^44^ This has been observed in the brain microvasculature of cerebral malaria patients, where free Hz crystals are observed next to the endothelial cells,^30^ suggesting that there is close contact of Hz released by iRBCs with the brain endothelium. Further, we also observed that purified *P. falciparum* Hz was able to disrupt the endothelial barrier of different cell types, including pulmonary endothelial cells, which may implicate Hz-bound biomolecules in the pathogenesis of other complications of severe malaria such as respiratory distress, where iRBCs are sequestered in other organs besides the brain.^38^ This is further supported by previous in vivo research where it was observed that *P. berghei* mutants with decreased Hz levels did not induce bone loss compared to the wild type control,^47^ highlighting the importance of Hz in malaria pathogenesis throughout organ systems other than the brain.

In line with previous reports that implicate proteins such as *P. falciparum* histones and HRPII in barrier disruption,^16–19^ we observed that treatment of *P. falciparum* Hz with proteases inhibits their HBMEC barrier-disrupting ability, indicating a role for proteins in the barrier disruption. Our findings also show that proteins within the iRBC lysate supernatant fraction, that are therefore not bound to Hz, do not present barrier disrupting activity at the physiological concentrations used in our assay.^33^ Previous reports have described in vitro barrier-disrupting activity with high concentrations of purified *P. falciparum*-iRBC soluble factors.^16–20^ However, it is possible that the concentrations required for endothelial disruption may be lower in the actual context of CM, when these biomolecules are bound to Hz and are therefore presented in high local concentrations to the endothelial cell surface.

Hz-bound proteins may also play a structural role, functioning as a physical link between the Hz crystal and the active biomolecules, such that when proteins are removed from the Hz, so are the active biomolecules. This may be supported by the SEM images of protease-treated Hz purified from iRBC lysate, which appears ‘naked’. Further work must be done to identify potential contributing roles of other types of biomolecules, such as nucleic acids or lipids.

The identification of Hz-bound proteins as mediators of endothelial disruption in an in vitro model of CM is an important step in understanding the pathogenesis of CM. Further work should be done to validate this mechanism in patients and identify the specific biomolecules that are contributing to the BBB disruption in CM. As there are currently no specific therapies for CM, finding treatments to protect the BBB from these specific biomolecules would drastically decrease the high morbidity and mortality rates of this fatal complication of malaria.

## METHODS

### Culture of endothelial cells

HBMECs from ScienCell were immortalized as previously described^14^ and used up to passage 18. Primary HBMECs from Cell Systems (ACBRI 376V) were used up to passage 13. HPMECs were immortalized as previously described^48^ and used up to passage 14. These three cell types were maintained in Endothelial Cell Media (ScienCell 1001) supplemented with 5% FBS (ScienCell 0025), 1% endothelial cell growth supplement (ScienCell 1052), and 1% penicillin/streptomycin solution (PS, ScienCell 0503). Experiments were performed in serum-free endothelial cell media supplemented only with 1% PS. HCMECs from PromoCell (C-12285) were used up to passage 8. HCMECs were maintained in Endothelial Cell Growth Media MV (PromoCell C-22120) supplemented with 5% FCS (PromoCell C-37310), 0.4% endothelial cell growth supplement with 90µg/mL heparin (PromoCell C-30120), 10ng/mL recombinant human epidermal growth factor (PromoCell C-30226), and 1µg/mL hydrocortisone (PromoCell C-31061). HCMEC experiments were performed in endothelial cell growth media MV (PromoCell) with no supplements. All cells were maintained at 37°C in 5% CO2, grown to ∼95% confluency then seeded for experiments, and were confirmed mycoplasma-free using the MycoAlert Mycoplasma Detection Kit (Lonza LT07-418).

### Culture, isolation, and lysate preparation of *Plasmodium falciparum*

*Plasmodium falciparum* 3D7 was used unless otherwise noted. Dd2 (MRA150), NF54 (MRA1000), and W2 (MRA157),^49^ were kindly provided by BEI resources (Manassas, VA). *P. falciparum* was maintained in human RBCs from healthy donors (Grifols, Vista, CA) at 5% hematocrit in RPMI 1640 (Corning 10-040-CV) supplemented with 25mM HEPES (Fisher Scientific BP310), 25mM sodium bicarbonate (Sigma-Aldrich S5761), 0.5mM hypoxanthine (Sigma-Aldrich H9377), 0.5% Albumax II (Thermo Fisher 11021045), and 10µg/mL gentamycin (Thermo Fisher 15750078). Mycoplasma-free parasite cultures were maintained at 37°C in a gas mixture of 5% O2, 5% CO2, and 90% N2, and were synchronized with 5% sorbitol (Sigma-Aldrich S1876) in water (Sigma W3500). Trophozoite- or schizont-stage iRBCs were isolated from synchronous cultures using magnetic columns (Miltenyi Biotec 130-042-901). Naturally ruptured iRBCs were generated by incubating 100µL aliquots of 200,000 isolated schizont iRBCs/µL overnight at 37°C in a flat-bottom, 96-well plate. iRBC lysate was generated by freeze-thawing isolated iRBCs 10x in liquid nitrogen and a 37°C water bath. Naturally ruptured iRBCs/iRBC lysate or the same number of control, donor-matched RBCs/RBC lysate were used for each experiment at a density of 8×10^6^ iRBCs/cm^2^. This concentration was described by Zuniga et al (2022)^33^ to correspond to two densely packed layers of iRBCs over the monolayer of HBMECs, which is representative of the accumulation of iRBCs in brain capillaries of patients with CM.^12^

### Hemozoin

#### Percoll purification

iRBC lysate or naturally ruptured iRBCs were carefully layered on a cold 10% Percoll solution (Sigma-Aldrich P4937 diluted in PBS (Fisher Scientific BP3994 in Milli-Q water))^35,36^ in a 15mL conical tube and spun at 3,000*g for 30min at 4°C with 0 acceleration and brake. Supernatant on top of the Percoll layer was removed and saved. The Percoll layer was aspirated, leaving only the Hz pellet. The pellet was collected and washed 2x in cold PBS at 21,130*g for 4min at 4°C. Hz was resuspended in colorless endothelial cell media (ScienCell 1001-b-prf) and was used fresh or frozen overnight at -80°C.

#### Magnet purification

A 1.5mL tube of iRBC lysate was attached to the top of a strong magnet (Applied Magnets NB041 or Miltenyi Biotec 130-090-976)^50^ for 45min to 1h, then the supernatant was removed. The Hz pellet was collected and washed 2x in cold PBS at 21,130*g for 4min at 4°C. Hz was resuspended in colorless media and was used fresh or frozen overnight at -80°C.

#### Hemozoin quantification

Samples were de-crystallized in a 96-well plate in a total volume of 200µL of a solution of 30mM sodium hydroxide (NaOH, Fisher Scientific S318) and 2% sodium dodecyl sulfate (SDS, Sigma-Aldrich L6026) in Milli-Q water overnight at room temperature (RT).^51–53^ Samples were quantified against a standard curve of synthetic hemozoin (InvivoGen tlrl-hz) reconstituted in 30mM NaOH + 2% SDS following measurement of absorbance at 405nm using the PerkinElmer VICTOR Nivo (PerkinElmer, Waltham, MA).

#### Synthetic hemozoin

Synthetic Hz was suspended in colorless media, aliquoted, quantified as above, and kept at 4°C. Synthetic Hz was mixed thoroughly with a micropipette before being diluted to the appropriate concentration.

### Measurement of barrier integrity

Barrier integrity was measured as electrical impedance by the xCELLigence RTCA DP (Agilent, Santa Clara, CA) at 37°C and in 5% CO2. 12,500-15,000 endothelial cells were seeded on collagen-coated (40µg/mL, Sigma-Aldrich C3867) RTCA PET E-Plate VIEW 16 plates (Agilent 300600880) and grown until the cells reached a plateau, after approximately 20-26h, with impedance measurements taken every 3h. The cells were washed with the serum-free media described above before the different experimental conditions were added to the cells. Impedance measurements were taken every 15min over 24h. The cell index value of the final measurement of the growth curve for each well was used to normalize the experimental cell index values throughout the 24h of the experiment.^33^

### Measurement of monolayer permeability

15,000 HBMECs were seeded on 0.4µm filter inserts in Transwell plates (Corning 3470) and grown for 24h. The media was aspirated from the wells and inserts and changed to the serum-free media described above. Media, the Percoll supernatant, or 18.88µg/cm^2^ purified Hz from iRBC lysate or synthetic Hz, were added to the monolayer of cells in the inserts for 24h. The media from the inserts was removed and the inserts were moved to new wells containing colorless media. 1mg/mL of 70,000 MW fluorescein-dextran (Thermo Fisher D1823) was added to the inserts for 20min at RT. The inserts were removed and the fluorescence intensity was measured at 485/535 nm. The fold change was calculated compared to the values from the HBMECs incubated with media.

### Scanning electron microscopy (SEM)

The samples were processed and imaged by the NYU Microscopy Laboratory. 12mm glass coverslips were cleaned with ethanol, coated with poly-l-lysine (Ted Pella Inc. 18026), then rinsed 3x with distilled water. 1-3 nanomoles of sample in distilled water were applied onto the glass coverslips in a 24-well plate. The Hz formed a thin layer on the glass surface following centrifugation at 3,000*g for 10min. Each well was incubated with 1mL of fixative (2.5% glutaraldehyde (Electron Microscopy Sciences 16019) with 0.1% malachite green (Thermo Fisher 611255000)^54,55^ in 0.1M sodium cacodylate buffer (Electron Microscopy Sciences 11654), pH 7.2) for 30min at RT, then washed 3x with water for 1min. The fixed samples were stained with 2% uranyl acetate in water for 30-60min at RT with slow shaking, then quickly rinsed in 50%; 70%; 90% ethanol, and washed 3x with 100% ethanol for 1min. Samples were critical point dried with the Tousimis Autosamdri-931 critical point dryer (Tousimis, Rockville, MD), then coated with gold/palladium using the Safematic CCU-010 SEM coating system (Rave Scientific, Somerset, NJ). Processed samples were imaged with the ZEISS Gemini300 FESEM (ZEISS, Oberkochen, Germany) using a secondary electron detector at 4.00kV with a working distance of 9.5mm.

### Imaging of Giemsa slides

Giemsa-stained (Sigma-Aldrich GS1L) slides were imaged with the AmScope MD500 camera (AmScope, Irvine, CA) attached to the Olympus BX41 microscope (Evident Scientific, Waltham, MA) and 100x oil objective (Olympus 1-UB235).

### HBMEC fluorescence microscopy

12,500 HBMECs were grown in collagen-coated, glass-bottom plates (Greiner Bio-One 655981) for 24h at 37°C then washed with serum-free media. Indicated conditions were added to the plate and incubated for the indicated time at 37°C. Cells were washed 2x with warm PBS, fixed with 4% paraformaldehyde (Santa Cruz Biotechnology CAS 30525-89-4) for 10min at RT, and washed 2x with PBS.

For assessment of endothelial disruption, cells were stained with phalloidin (1:500 in PBS, Thermo Fisher A12381) for 15min at RT, washed 3x with PBS, stained with Hoechst (1:10,000 in Milli-Q water, Thermo Scientific H3570) for 5min at RT, and washed 3x with PBS. The cells were imaged with the Olympus IX70 inverted microscope (Evident Scientific, Waltham, MA) using a 60x/1.4 objective (Olympus 1-UB933) and MetaMorph Advanced software version 7.6.5.0 (Molecular Devices, San Jose, CA).

For Hz uptake, HBMECs were permeabilized with 0.25% Triton X-100 (Sigma-Aldrich T9284) for 10min at RT, then washed 3x with PBS, incubated with blocking buffer (10% goat serum, 1% BSA (Sigma-Aldrich A3912), 100mM glycine (Sigma-Aldrich 410225), and 0.05% sodium azide (Sigma-Aldrich S832) in PBS) for 30min at RT, and incubated with α-LAMP1 antibody (1:100 in PBS + 5% blocking buffer, Cell Signaling Technology 9091T) for 1h at RT. The cells were washed 2x with PBS then incubated with α-rabbit antibody (1:500 in PBS + 5% blocking buffer, Thermo Fisher A-11008) for 1h at RT. Cells were washed 2x with PBS and stained with Hoechst. The cells were imaged with a ZEISS AxioObserver with a 63x/1.4 lens, standard narrow pass fluorescence filters with a Hg arc lamp, an Axiocam 503 monochrome CCD camera, and Zen Blue 2.6 software (ZEISS, Oberkochen, Germany).

To ensure that all Hz particles are visualized in our experimental conditions (**Figure 4**), we imaged the same concentrations of iRBC lysate, Hz from iRBC lysate, and synthetic Hz without cells, finding that the dark Hz crystals in the brightfield images correspond to the white spots under crossed polarization (**Figure S9**), demonstrating that the Hz in the iRBC lysate and Hz purified from the iRBC lysate would be detected in the HBMECs if they were internalized like the synthetic Hz.

### Experimental treatments of iRBCs or Hz

The iRBC lysate was spun at 16,100*g for 3min to separate the supernatant and pellet, and pellet was resuspended in colorless media. iRBC lysate was heated at the indicated temperature and times then cooled on ice for 5min. iRBC lysate tubes were sonicated one at a time in a cup horn containing ice water for 5min at 100% power (Qsonica Sonicators Q500). iRBC lysate was incubated with different volumes of either media, hydrochloric acid (HCl, Fisher Scientific A144) or NaOH for 30min at 37°C, then the pH was measured with pH-indicator strips. TURBO DNase (Thermo Scientific AM2238) at 40U with 1X DNase Buffer, RNase A (Thermo Fisher EN0531) at 100µg/mL, trypsin (Corning 25-053-CI) at 57.6µg/mL or media was incubated with the indicated stimuli for 30min at 37°C. iRBC lysate or Hz were incubated with Proteinase K (Novagen 71049-3) or media at 10µg/mL for 30min before spinning at 16,100*g for 3min at RT, removing the supernatant, and reconstituting the pellet in colorless media.

### Measurement of DNA and RNA

10µL of sample were incubated with 0.2µL/mL SYBR Green (Thermo Fisher S7563) for DNA, 500nM SYTO RNASelect (Thermo Scientific S32703) for RNA, or PBS for unstained in a 96-well plate at a total volume of 100µL for 1h at RT, shaking. Fluorescence was measured at excitation of 485nm and emission at 530nm. The fluorescent signal for each unstained sample was subtracted from the corresponding stained sample. These values were then corrected by subtracting a blank stained with either SYBR Green or SYTO RNASelect, then were normalized to the untreated iRBC lysate values.

### Silver Stain

iRBC lysate and Hz from iRBCL were generated as above. Protein was measured with the Pierce 660nm Protein Assay Reagent (Thermo Fisher 22660). The protein samples were then diluted in Laemmli Sample Buffer (Bio-Rad 1610747) supplemented with 10% β-mercaptoethanol (Sigma-Aldrich M7522), boiled at 95°C for 5min, cooled on ice, and spun down at 16,100*g to remove remaining Hz. 200ng of protein and the Precision Plus Protein Standard (Bio-Rad #161-0374) were loaded in a 12% polyacrylamide gel (Bio-Rad 4561045) and run in Tris-Glycine SDS Buffer (Thermo Fisher 28362) on ice 2h at 100V. The gel was silver stained using the Pierce Silver Stain Kit (Thermo Fisher 24612).

### Proteomics

Proteomics was performed by the NYU Proteomics Laboratory. 70µL of Hz from iRBC lysate was diluted 1:1 with 10% SDS, reduced with 2µL of 0.2M dithiothreitol (DTT) at 57°C for 1h, alkylated with 2µL of 0.5M iodoacetamide for 45min at RT in the dark, then quenched with 4µL 0.2M DTT. Sample was acidified with 12% phosphoric acid to a pH < 2 then diluted 1:6 in 100mM triethylammonium bicarbonate (TEAB) in 90% methanol then loaded onto an S-Trap column (ProtiFi, Fairport, NY) in 150µL increments using a microcentrifuge. Columns were washed 5x with 100mM TEAB in 90% methanol then spun to remove any remaining buffer. The samples were digested on the S-Trap column with 2µg of trypsin in 100mM ammonium bicarbonate at 47°C for 2h. Peptides were eluted from the column with 40µL of 50mM TEAB and 100mM ammonium bicarbonate, then 40µL of 0.5% acetic acid, then 40µL of 50% acetonitrile into the same tube. The organic solvent was removed by vacuum drying the sample, and the sample was reconstituted in 0.5% acetic acid.

1/3 of the sample was analyzed on an EASY-nLC 1000 coupled to a Q Exactive Orbitrap mass spectrometer (Thermo Fisher Scientific, Waltham, MA). The peptides were gradient eluted with solvent A being 2% acetonitrile in 0.5% acetic acid and solvent B being 80% acetonitrile in 0.5% acetic acid. The gradient was held for 5min at 5% solvent B, ramped in 120min to 35% solvent B, in 10min to 45% solvent B, and in another 10min to 100% solvent B. High resolution full mass spectrometry (MS) spectra were acquired with a resolution of 70,000, an automatic gain control (AGC) target of 10^6^, with a maximum ion time of 120ms, and scan range of 400-1500 m/z. The top 20 higher-energy collisional dissociation MS/MS spectra were collected using the following instrument parameters: resolution of 17500, AGC target of 5×10^4^, maximum ion time of 120ms, one microscan, 2 m/z isolation window, first mass 150, and nominal collision energy of 27.

The resulting MS/MS spectra were searched against the *P. falciparum* 3D7 database, the Human database, and the universal contaminant protein (UCP) database using Sequest within Proteome Discoverer 2.5 (Thermo Fisher, Waltham, MA). Search parameters were as follows: 10ppm MS1 mass tolerance, 0.02Da MS2 mass tolerance, 2 max missed cleavages, minimum peptide length of 6 amino acids. Allowed modifications were oxidation on methionine and tryptophan (dynamic), deamidation on asparagine (dynamic), and carbamidomethyl on cystine (fixed). A target-decoy approach with a 1% FDR cutoff was used to filter the results. 2 peptides per protein were required to make the protein identification.

### Statistics

Statistical analyses were performed with GraphPad Prism 10 (GraphPad Software, Boston, MA). Normality was first determined with the Shapiro-Wilke test. Data of normal distribution was analyzed by one-way ANOVA with Tukey’s multiple comparisons test or unpaired t test. Data of non-normal distribution was analyzed by Kruskal-Wallis test with Dunn’s multiple comparisons test or Mann-Whitney test. The statistical test used for each figure is denoted in the figure legend.

## AUTHOR CONTRIBUTIONS

K.A.C. and A.R designed and supervised the project. K.A.C. carried out experiments. I.C. and B.U. were responsible for the proteomics. A.L. and J.S. were responsible for the electron microscopy. M.C. assisted with cross polarization and fluorescence microscopy. K.A.C. and A.R. were responsible for manuscript preparation and revision. All authors read and approved the final manuscript.

## ACKNOWLEDGEMENTS

The following reagents were kindly provided by BEI Resources, NIAID, NIH: *Plasmodium falciparum* Strain Dd2 (MRA-150, contributed by David Walliker), Strain NF54 Patient Line E (MRA-1000, contributed by Megan G. Dowler), and Strain W2 (MRA-157, contributed by Dennis E. Kyle). The primary human cardiac microvascular endothelial cells (HCMECs) were kindly provided by Kenneth Stapleford at the NYU School of Medicine. This work was supported by NIH/NIAID R01AI181219, R01HL150145, and R01NS105910 to A.R., and T32AI007180 to K.A.C. The NYU Microscopy Laboratory and NYU Proteomics Laboratory are partially supported by NYU Cancer Center Support Grant NIH/NCI P30CA016087, the Gemini300 FESEM was supported by NIH S10OD019974, and the Orbitrap Fusion Lumos Tribrid mass spectrometer was supported by the NIH Shared Instrumentation Grant 1S10OD010582-01A1.

