## Supplemental Figures for "*Plasmodium falciparum* hemozoin-associated biomolecules induce brain endothelial cell barrier disruption in an in vitro model of cerebral malaria"

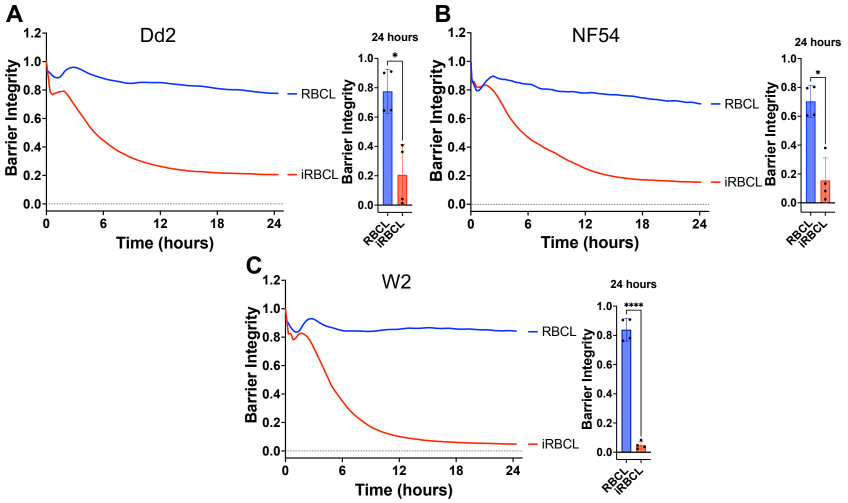


**Figure S1: iRBC lysate from different *P. falciparum* strains disrupt the junctions of HBMECs.** RBC lysate (RBCL) or Dd2- (**A**), NF54- (**B**), or W2- (**C**) iRBC lysate (iRBCL) were added to a monolayer HBMECs. Barrier integrity of HBMEC monolayers was measured by xCELLigence as impedance every 15min over 24h. Bars represent barrier integrity at 24h. Results represent the average from 2 independent experiments performed in duplicate with standard deviations. Statistical significance was determined by Mann-Whitney test (**A-B**) or unpaired t test (**C**) where * = $P < 0.05$ and **** = $P < 0.0001$. (**A, C**) RBCL line was smoothed using 2$^{nd}$ order smoothing with 6 neighbors using GraphPad Prism 10.

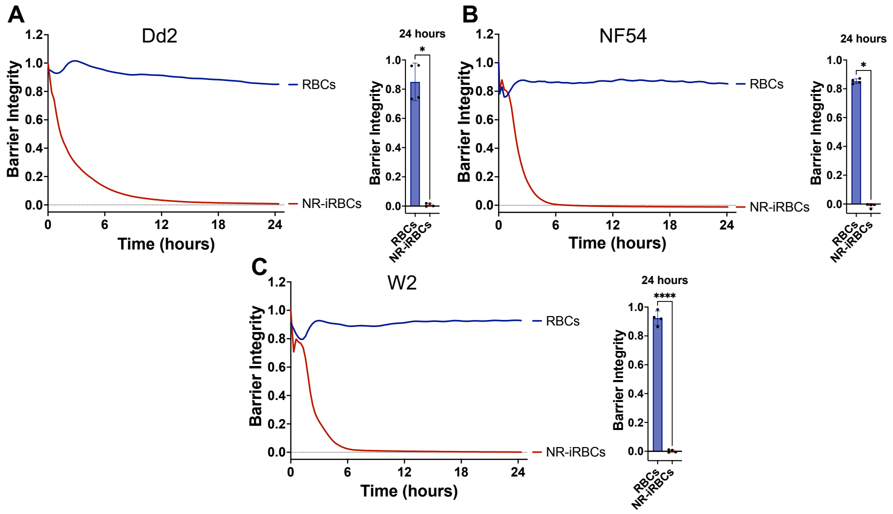


**Figure S2: Naturally ruptured iRBCs from different *P. falciparum* strains disrupt the junctions of HBMECs.** RBCs or Dd2- (**A**), NF54- (**B**), or W2- (**C**) naturally ruptured iRBCs (NR-iRBCs) were added to a monolayer HBMECs. Barrier integrity of HBMEC monolayers was measured by xCELLigence as impedance every 15min over 24h. Bars represent barrier integrity at 24h. Results represent the average from 2 independent experiments performed in duplicate with standard deviations. Statistical significance was determined by Mann-Whitney test (**A-B**) or unpaired t test (**C**) where * = $P < 0.05$ and **** = $P < 0.0001$. (**A, C**) RBCs line was smoothed using 2$^{nd}$ order smoothing with 6 neighbors using GraphPad Prism 10.

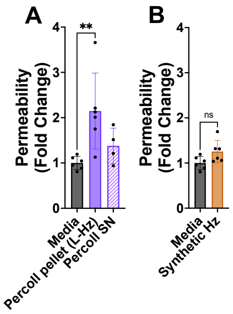


**Figure S3: Purified Hz from iRBC lysates, but not synthetic Hz, induces permeability of HBMEC monolayers.** Schizont-stage iRBC lysate was spun over a 10% Percoll solution to obtain pellet and SN. (**A**) The Percoll supernatant (SN), the Percoll pellet containing Hz (18.88μg/cm$^2$), or (**B**) synthetic Hz at the same concentration were added to a monolayer of HBMECs for 24h before measuring permeability. The leakage of 70,000 MW fluorescein-dextran is expressed as fold change compared to the media control. Results represent the average from 2+ independent experiments performed in duplicate with standard deviations. Statistical significance was determined by one-way ANOVA with Tukey's multiple comparisons test where ** = $P$ < 0.01.

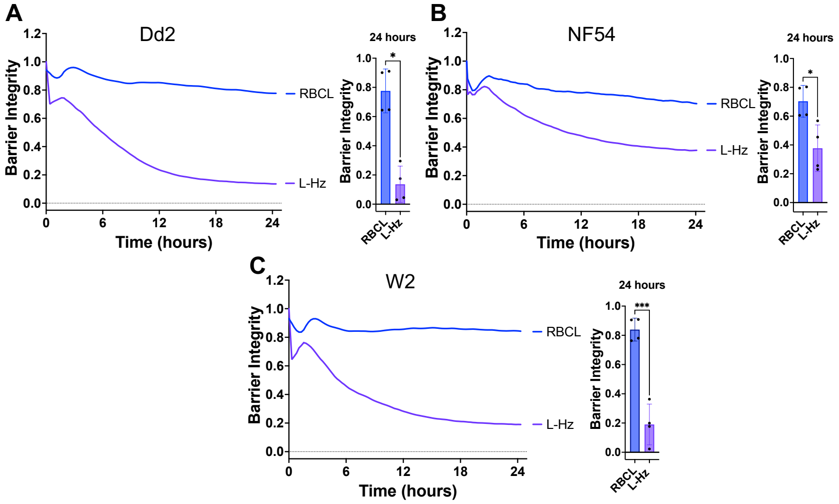


**Figure S4: Hz from iRBC lysates of different *P. falciparum* strains disrupt the junctions of HBMECs.** RBC lysate (RBCL) or Dd2- (**A**), NF54- (**B**), or W2- (**C**) Hz from iRBC lysate (L-Hz; 18.88μg/cm$^2$) were added to a monolayer HBMECs. Barrier integrity of HBMEC monolayers was measured by xCELLigence as impedance every 15min over 24h. Bars represent barrier integrity at 24h. Results represent the average from 2 independent experiments performed in duplicate with standard deviations. Statistical significance was determined by Mann-Whitney test (**A-B**) or unpaired t test (**C**) where ** = $P$ < 0.01 and **** = $P$ < 0.0001. (**A, C**) RBCL lines were smoothed using 2$^{nd}$ order smoothing with 6 neighbors using GraphPad Prism 10.

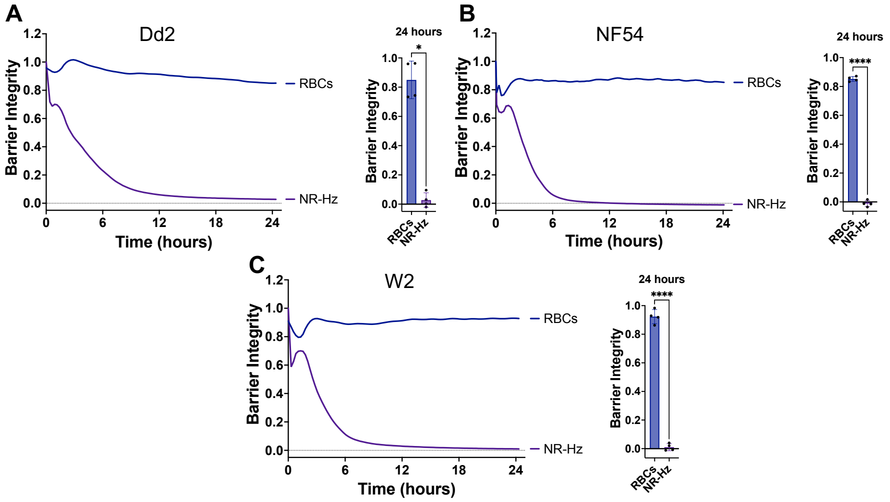


**Figure S5: Hz from naturally ruptured iRBCs of different *P. falciparum* strains disrupt the junctions of HBMECs.** RBCs or Dd2- (**A**), NF54- (**B**), or W2- (**C**) Hz from naturally ruptured iRBCs (NR-Hz; 18.88μg/cm$^2$) were added to a monolayer HBMECs. Barrier integrity of HBMEC monolayers was measured by xCELLigence as impedance every 15min over 24h. Bars represent barrier integrity at 24h. Results represent the average from 2 independent experiments performed in duplicate with standard deviations. Statistical significance was determined by Mann-Whitney test (**A**) or unpaired t test (**B-C**) where ** = $P < 0.01$ and **** = $P < 0.0001$. (**A, C**) RBCs line was smoothed using 2$^{nd}$ order smoothing with 6 neighbors using GraphPad Prism 10.

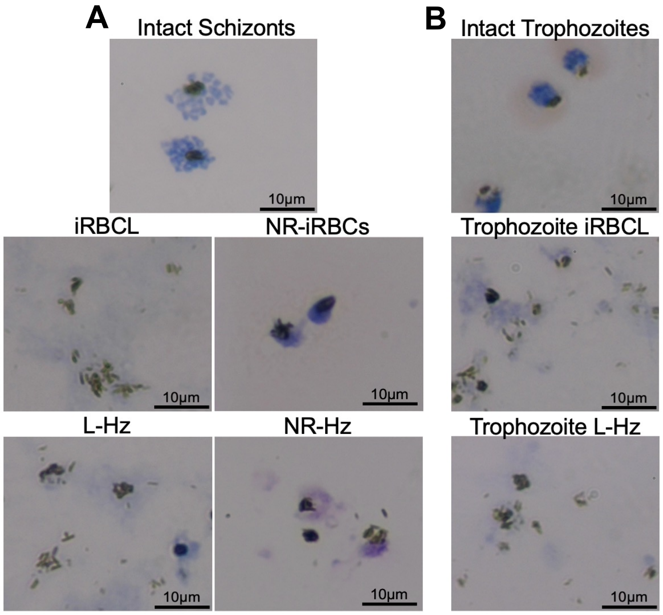


**Figure S6: Giemsa-stained images of schizont and trophozoite iRBCs and Hz.** Representative images of schizont-stage (**A**) or trophozoite-stage (**B**) iRBCs that were imaged before and after rupture by lysing (iRBCL) or natural rupture (NR-iRBCs), and after Percoll-purification of Hz from iRBCL (L-Hz) or NR-iRBCs (NR-Hz).

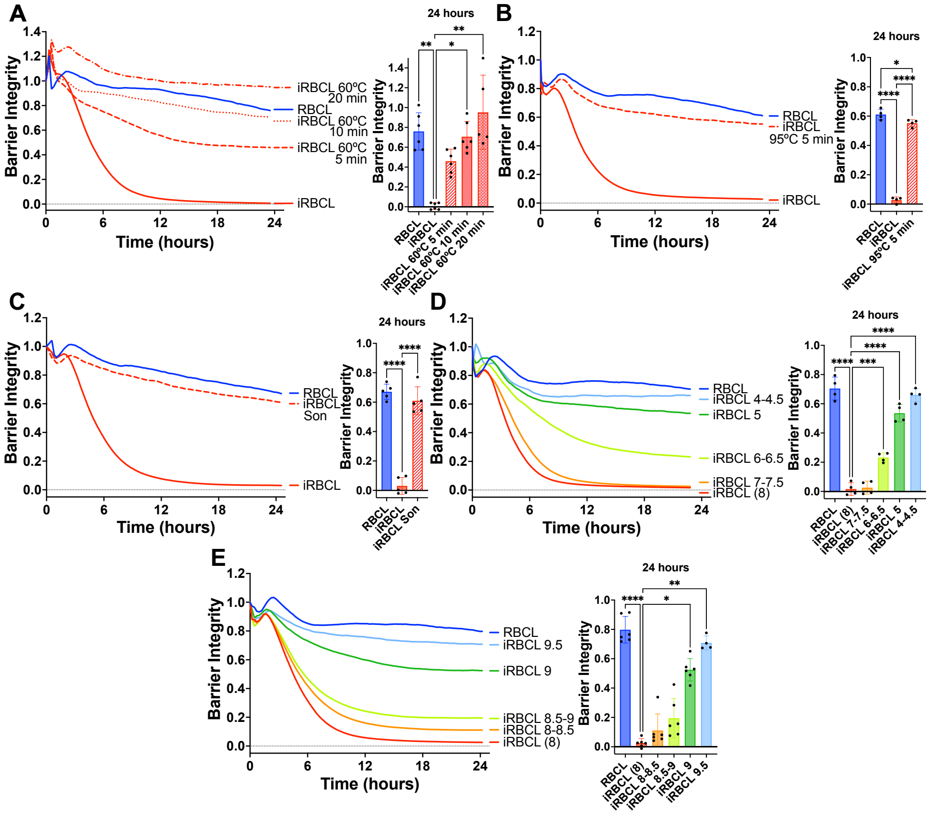


**Figure S7: Physical and chemical changes to the iRBC lysate cause the loss of disruptive ability**. iRBC lysate (iRBCL) was heated at 60ºC for the indicated times (**A**) or 95ºC for 5min (**B**) before adding to a monolayer of HBMECs along with RBC lysate (RBCL) as a control. (**C**) iRBCL was sonicated (Son) on ice at max power for 5min before adding to a monolayer of HBMECs along with RBCL as a control. The pH of the iRBCL was brought from its natural pH of 8.0 down to the indicated pH with hydrochloric acid (**D**) or up to the indicated pH with sodium hydroxide (**E**), then incubated at 37ºC for 30min before adding to a monolayer of HBMECs along with RBCL as a control. Barrier integrity of HBMEC monolayers was measured by xCELLigence as impedance every 15min over 24h. Bars represent barrier integrity at 24h. Results represent the average from 2+ independent experiments performed in duplicate with standard deviations. Statistical significance was determined by Kruskal-Wallis test with Dunn's multiple comparisons test (**A, E**) and one-way ANOVA with Tukey's multiple comparisons test (**B, C, D**) where * = $P < 0.05$, ** = $P < 0.01$, *** = $P < 0.001$, and **** = $P < 0.0001$.

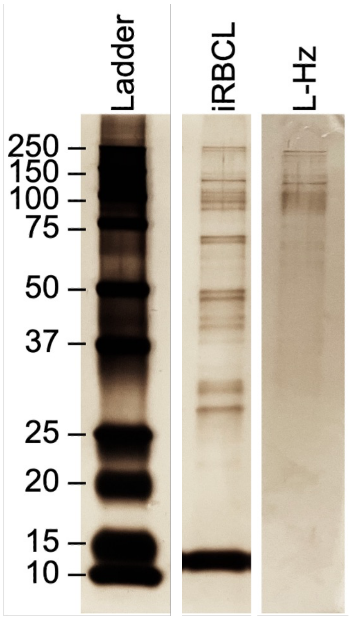


**Figure S8: Many *P. falciparum* iRBC proteins are bound to Hz.** SDS-PAGE separation of 200ng of protein iRBC lysate (iRBCL) and Hz from iRBCL (L-Hz) was silver stained to visualize proteins. Representative image of 3 experiments. Gel image was cropped to remove unrelated conditions; cuts are indicated by spaces between ladder and iRBCL, and iRBCL and L-Hz.

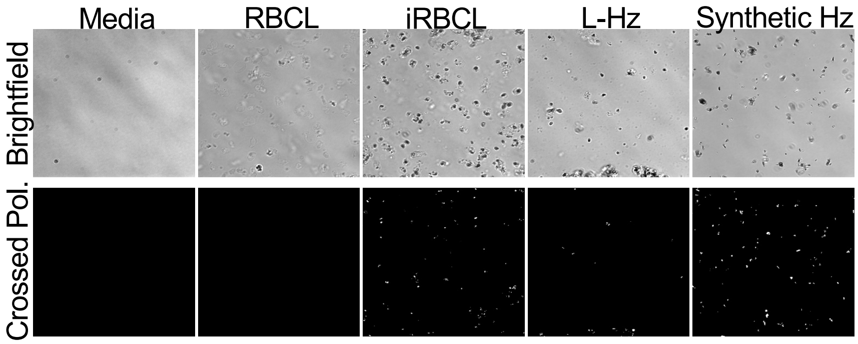


**Figure S9: Hz can be visualized under crossed polarization.** RBC lysate (RBCL), iRBC lysate (iRBCL), Hz from iRBCL (L-Hz), or synthetic Hz (concentration matched to Hz in iRBCL)) were added to a 96-well plate without cells and were imaged under brightfield and crossed polarization (Crossed Pol.).
